# Overexpressed Nup88 stabilized through interaction with Nup62 promotes NFκB dependent pathways in cancer

**DOI:** 10.1101/2020.04.27.063057

**Authors:** Usha Singh, Atul Samaiya, Ram Kumar Mishra

## Abstract

Nuclear pores control nucleo-cytoplasmic trafficking and directly or indirectly regulate vital cellular processes. Nup88, important for Crm1 mediated nuclear export process, is overexpressed in many cancers. A positive correlation exists between progressive stages of cancer and Nup88 expression. However, links between Nup88 overexpression and head and neck cancer are insignificant, and mechanistic details are non-existent. Here, we report that Nup88 exhibits positive correlation in head and neck cancer in addition to elevated Nup62 levels. We demonstrate that Nup88 interacts with Nup62 in a cell-cycle and glycosylation independent manner. The overexpression of Nup88 or Nup62 imparts proliferation and migration advantages to cells. We further report that the interaction with Nup62 stabilizes Nup88 by inhibiting proteasome-mediated degradation of overexpressed Nup88. Overexpressed Nup88 is stabile and partly inside the nucleus and can interact with NFκB (p65). Nup88 overexpression induces proliferative and inflammatory responses downstream of p65. Altogether, we suggest that simultaneous overexpression of Nup62 and Nup88 in head and neck cancer stabilizes overexpressed Nup88. Stable Nup88 interacts with p65 and induces inflammatory, proliferative, and migratory advantages to cells, which perhaps is the underlying mechanism driving tumorigenic transformations.

## Introduction

Nucleoporins (Nups) are the constituent protein of the megadalton assemblies called nuclear pores. ~30 different Nups form biochemically distinct and stable sub-complexes with precise perinuclear localization. During interphase, Nups localize to the nuclear membrane and mediate nucleo-cytoplasmic transport, but, in mitosis, subcomplexes disassemble, and some of them localize to chromatin and regulate mitotic spindle assembly, microtubule dynamics, chromosome segregation ^1–3^. Dynamic localization and functional roles of nucleoporins make them vital in affecting cellular functions and inducing diseases, like cancer.

Nup88, in complex with Nup214, is found at the cytoplasmic face of the nuclear pores ^4^. Point mutations and expression level changes in nucleoporins have links with occurrences and progression of cancer ^5^. Nup88 mRNA and protein levels were reported to be enhanced in human ovarian tumors ^6^. Elevated Nup88 levels were found in several cancers irrespective of their type, degree of differentiation, or site of occurrence ^7^. Moreover, Nup88 levels exhibit a positive correlation with progressive stages of cancer ^8^. Reports of a strong Nup88 reactivity at the periphery and edges of the advanced stage tumors exist ^7^. CAN (Nup214), a proto-oncogene linked with myeloid leukemiogenesis ^9^, forms a complex with Nup88 and regulates CRM1 mediated nuclear export of macromolecules ^10^. Nonetheless, Nup214 is not co-expressed in Nup88 overexpressing cancers ^11^. Overexpression of Nup88 induced multinucleated phenotypes, and a multipolar spindle phenotype when depleted. Interestingly, Nup214 co-expression in Nup88 overexpressing cells ameliorated above phenotypes, highlighting the importance of free levels of Nup88 and its complexation with Nup214 in cellular homeostasis ^12^. Moreover, overexpression of Nup88 sequestered Nup98-Rae1 away from APC/C complex triggering early degradation of PLK1 that induced aneuploidy and tumorigenesis ^13^. Also, the interaction of Nup88 with Vimentin affects the Vimentin organization resulting in multinucleated cells and aneuploidy ^14^.

Nup159, the yeast ortholog of Nup214, is mono-ubiquitinated and affects the cell-cycle progression and aneuploidy ^15^. In yeast, Nup88 interacted with Nup62 through the helical domain ^16^ and mutations in Nup62 affected the mRNA export ^17^. Nup62 glycosylation is an important determinant of Nup88 stability ^18^, and the ubiquitination of Nup88 and Nup62 affects their stability ^15^. Moreover, Nup88 interacted with Nup98 ^19^, Nup358 ^20^, and lamin A ^21^ to modulate various processes at the nuclear periphery. Nup62 overexpression is reported from the prostate, and ovarian cancers ^22,23^, and ROCK1 dependent Nup62 phosphorylation induces p63 nuclear localization and cell proliferation ^24^. Multiple physiological roles attributed to Nup88 ^25^ do not provide a clear picture, and the availability of limited information about the Nup88 and Nup62 expression level changes in head and neck cancer ^11,24^ impedes our understanding of the process. Since Nup88 and Nup62 form stable complex and their expression levels show alterations in different cancers, we have probed how the expression and interactions of Nup88 and Nup62 correlate with head and neck cancers, common cancer in India.

Here, we report that head and neck cancer tissues display enhancement in the Nup88 and Nup62 mRNA and protein levels. Overexpression of Nup88 and Nup62 shows a positive correlation with oral cancer. Overexpression of Nup88 imparts proliferation and migration potential in cells. The conserved Nup88-Nup62 interaction is through their respective carboxy-terminal regions and is independent of cell cycle dynamics and glycosylation status of Nup62. Nup62 co-overexpression in cancer cell lines, primarily stabilizes and prevents ubiquitination mediated degradation of Nup88. Stabilized Nup88 interacts efficiently with NFκB and affects proliferation, inflammation, and anti-apoptosis responses downstream of NFκB signalling to promote tumorigenic growth.

## Results

### Nup88 and Nup62 are overexpressed in head and neck cancer

We asked how the Nup88 expression level changes correlate with head and neck (oral) cancer. We started with probing of Nup88 expression levels in oral cancer tissues and used Nup62 as another nucleoporin control. When compared with normal tissues obtained from the vicinity of the tumor, Nup88 levels were higher in tumor tissues, and coincidently the Nup62 levels were also high (Fig. 1a, n=4). A variation in the control gene (GAPDH) levels is evident, but the ratiometric analysis of Nup62 or Nup88 with GAPDH indicates that both Nups are significantly overexpressed in oral cancer tissues (Fig. 1b, c, n=4). We further asked if the increased Nup88 and Nup62 transcript levels are behind their increased protein levels. Using Rps16 as a loading control, we determined Nup62 and Nup88 levels and noticed a distinct change (1.2 and 1.6 fold respectively) in their mRNA levels (Fig. 1d, e, n=7). Analysis of Nup62 protein levels in more oral cancer tissues suggests an invariably higher level (~2 fold) in tumor tissues than in normal tissues (Fig. 1f, S1a n=10). Next, we analyzed the expression of Nup62 and Nup88 using the Oncomine database ^26^ in the Ginos Head and neck cancer statistics ^27^. The analysis revealed a 2.1 and 1.15 fold increase, respectively, in the transcript levels of Nup62 and Nup88 (Fig. S1b). We further asked if Nup62 upregulation is specific to oral cancer or a more general phenotype of cancer. To analyze this, we checked the expression of Nup62 and Nup88 in a collection of cancer datasets available at MiPanda. In cancer *vs* normal tissue data analysis, we found Nup62 levels to be upregulated along with Nup88 in all different types of cancer selected (lung, breast, stomach, head and neck) for the analysis (Fig 1g, h) We compared the expression of Nup88 and Nup62 in normal tissues, primary cancer tissues, metastasis, and cancer cell lines of head and neck carcinoma at MiPanda database. Nup88 expression was found to be significantly more in cancer tissues in comparison with normal tissues (Fig. 1i). The observation is in accordance with the known literature for Nup88. However, the Nup62 overexpression noted to be significantly high in cancer tissue is a novel observation of this study (Lower panel, Fig. 1i). We also, analyzed the co-expression of Nup62 and Nup88 in oral cancers in a collection of available datasets at MiPanda ^28^. The Nup62 and Nup88 transcript levels are optimal in normal tissues, but significant overexpression of both these transcripts was common in primary tumors. (Fig. S1c). Next, we asked if there is any correlation between Nup62 and Nup88 expression levels and sub-stages of oral cancer. Using Cancer RNA-Seq Nexus (CRN) ^29^, we observed an enhancement in Nup62 levels with the progressive stages of cancer (Fig. S1d). We asked how the expression level changes in Nup62 and Nup88 correlate with patient survival. The Kaplan-Meier survival curve generated using Onco-Lnc ^30^ suggested no significant change in survival upon Nup62 overexpression (Fig. S1e), but the Nup88 has a definite decrease in the overall survival (Fig. 1j).

**Fig. 1:**
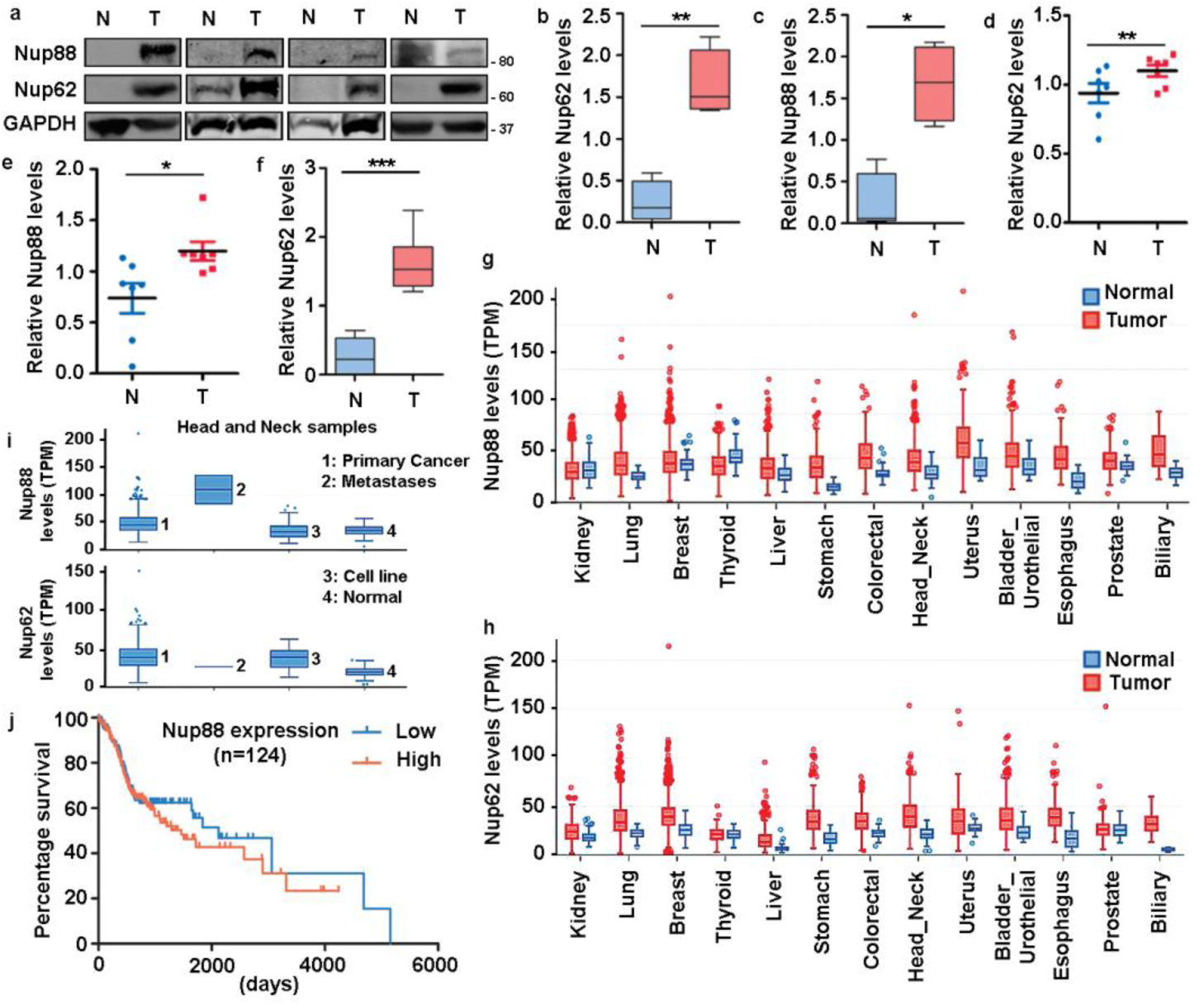
Nup88 and Nup62 are overexpressed in oral cancer. (a) Western blot analysis of lysates prepared from oral cancer tissues using antibodies against Nup88, Nup62, and GAPDH. (b, c) Quantification of Nup88 and Nup62 band intensities from normal and tumor tissues, shown in (a). Values on the y-axis represent data obtained by normalizing with the GAPDH band intensity. The asterisk indicates statistical significance p<0.05. (d, e) Graphs indicating fold changes in mRNA in the Nup62 and Nup88 levels, respectively, in oral cancer tissue samples (n=7). The values were obtained by normalizing to Rps16 levels. (f) Nup62 protein levels detection in oral cancer tissues (n=10). The values were obtained, as described in (b). (N=Normal adjacent tissues, T=Tumor). The asterisk represents the significance value. (Student’s t-test - paired t-test) *P < 0.05, **P < 0.01 and ***P < 0.001. (g) Nup88 (h) Nup62 overexpression in different cancers analyzed through MiPanda database. The boxplots were downloaded from MiPanda and used for the representation. (i) Nup88 and Nup62 expression in TCGA head and neck statistics analyzed in Mi-Panda. TPM (transcript per million). (j) Kaplan-Meier survival curve for Nup88 using OncoLnc on oral cancer TCGA data.

### Overexpression of Nup88 and Nup62 can induce tumorigenic transformations

Nup62 and Nup88 overexpression in cancer tissues and those analyzed from publicly available datasets hint that Nup62 overexpression can phenocopy the observations made with Nup88 overexpression. In a cell culture setup, we asked how Nup62 and Nup88 levels contribute to the vital characteristics like increased proliferation, migration, and loss of contact inhibition exhibited by cancerous tissues. Observations made through the MTT assays on MCF7a cells expressing GFP alone (Vector) or GFP-Nup62 or GFP-Nup88 full-length indicate a 4-7 fold increase in cell viability in GFP-Nup62 or GFP-Nup88 expressing cells as compared to GFP expression, suggesting an increased metabolic activity (Fig. 2a). We employed the wound healing experiment to assess growth and migration properties and observed that the wound healing was nearly complete by 72h post wounding (hpw) in Nup62 or Nup88 overexpressing cells, but it healed to only ~50% in control cells (Fig. 2b, c). Using colony-forming assay (CFA), we assessed the loss of contact inhibition property in MCF7a cells expressing GFP alone (Vector) or Nup62 or Nup88. As compared to GFP control, a significant difference in colony size was noticed in Nup62 or Nup88 expressing cells (Fig. 2d, e and Table S1). In a similar set up when Nup62 and Nup88 levels were reduced using shRNA, the number of colonies formed and their area significantly reduced when compared with control knockdown (Fig. 2f, g and Table S2). Careful analysis of these observations indicates that Nup88 overexpression has a greater effect on the tumorigenic phenotypes than Nup62 overexpression itself.

**Fig. 2:**
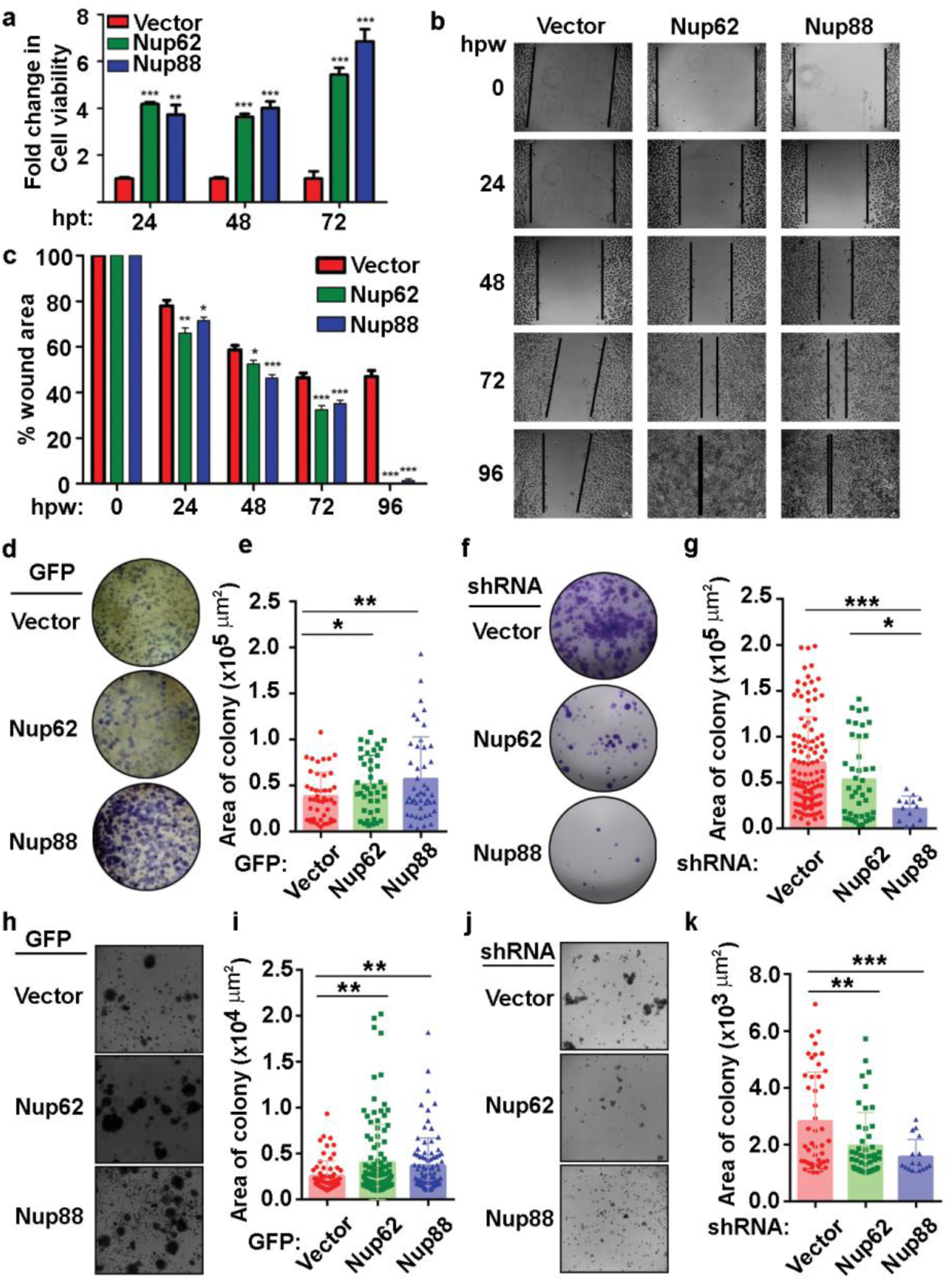
Overexpression of Nup88 and Nup62 induces tumorigenic transformations. (a) Fold change in cell viability assessed in GFP (Control), GFP-Nup62, and GFP-Nup88 transfected MCF-7a cells by MTT assay. The increase in cell proliferation was measured by making GFP (Control) as 100% at each time point, and relative fold change was calculated. (b) Representative image of wound healing assay in MCF-7a cells expressing GFP, GFP-Nup62, and GFP-Nup88 at different time points. The wound area lies within the space between two black lines in each panel. The scale bar is 500µm and the images were acquired at magnification of 10X under inverted microscope (c) Quantification of the closure of the wound area using TScratch software. (d) Colony formation assay in MCF-7a cells overexpressing GFP, GFP-Nup62, and GFP-Nup88. (e) Quantification of the area of colonies using Image-J software. (f) Colony formation assay with MCF-7a cells transfected with shRNA for Control, Nup62, and Nup88. (g) Quantification of the same using Image-J software. (h) Representative image of soft agar assay in MCF7a cells expressing GFP, GFP-Nup62, and GFP-Nup88. (i) Quantification of the same using Image-J software. (j) Soft agar assay with transfected cells as in (f). (k) Quantification of the same using Image-J software. Images are a representative from at least n=3 repeat experiments. Error bars show mean values ± SEM. Asterisks indicate statistical significance (Student’s t-test) *P < 0.05, **P < 0.01 and ***P < 0.001.

Further, using the soft agar assay, we asked if the expression of GFP or GFP-Nup62 or GFP-Nup88 in MCF7a cells (Fig. S2) can impart growth potential. As compared to vector control, an increase in colony size and their number was observed when Nup62 or Nup88 was overexpressed (Fig2. h, i, and Table S3). In similar soft agar assays, when shRNA mediated knockdown depleted Nup62 or Nup88 levels, a significant decrease in the number and colony size was observed as compared to vector control (Fig. 2j, k and Table S4). Our *in cellulo* observations with Nup62 and Nup88 expression level changes indicate that both Nup62 and Nup88 can induce tumorigenic transformations. Importantly, Nup88 appears to be a stronger and direct factor in the process.

### Nup88 and Nup62 engage in a conserved interaction through their carboxy termini

Next, we asked if Nup88 and Nup62 can physically interact like in yeast to mediate the Nup88 overexpression dependent cancer phenotypes. Using anti-Nup62 antibodies, we performed immunoprecipitation (IP) from HEK-293T and MCF7a cell lysates. Immunoblotting with Nup88 antibodies identified a strong band for Nup88 (Fig. 3a, b). Secondary structure prediction analysis on the Nup88 sequence (Uniprot ID: Q99567) suggests a much smaller coiled-coil domain than used in earlier studies. We generated Nup88 constructs which contain only the coiled-coil domain (Nup88C) and the one lacking the coiled-coil domain (Nup88ΔC) (Fig. 3c). In a pulldown experiment, recombinant GST-Nup88C purified from *E. coli* or GFP-Nup88 and GFP-Nup88C enriched on GFP binding protein (GBP) beads from mammalian cells efficiently pulled down endogenous Nup62 from HEK293T cell lysates (Fig. 3d, e). Accordingly, Nup88ΔC was unable to pull down Nup62 from HEK293T cell lysates (Fig. 3e). Further, we asked if the C-terminal alpha-helical region of Nup62 is involved in conserved interaction with Nup88. We generated the Nup62 truncations (Fig. 3f) as described elsewhere ^31^, and GSH-beads coated with GST-Nup62 truncations (N1, C1, and C2) were incubated with cell lysates expressing either GFP-Nup88, GFP-Nup88C or GFP-Nup88ΔC. The C-terminal alpha-helical region bearing truncations Nup62-C1 and Nup62-C2 pulled down Nup88 and Nup88C but not the Nup88ΔC (Fig. 3g). Thus the minimal alpha-helical region of Nup62 retained in the Nup62-C1 construct is required for Nup88-Nup62 interaction. In a reciprocal pulldown, GST-Nup88C was incubated with mammalian cell lysates expressing GFP-Nup62, Nup62ΔC1, or Nup62-C1. Nup62 and Nup62-C1 were pulled down efficiently on GST-Nup88C but not the Nup62ΔC1 (Fig. 3h). Using the yeast two-hybrid system, we further established that the minimal coiled-coil regions of Nup88 and Nup62 are sufficient to mediate the Nup88-Nup62 interaction (Fig. 3i). Importantly, Nup62-C1 does not interact with any random coiled-coil domain. The coiled-coil domain of intermediate filament binding protein Periplakin (PPL-C), does not pull down endogenous Nup62 and GFP-Nup62-C1 from HEK cell lysates, whereas Nup88C pulls down both these proteins efficiently (Fig. S3).

**Fig. 3:**
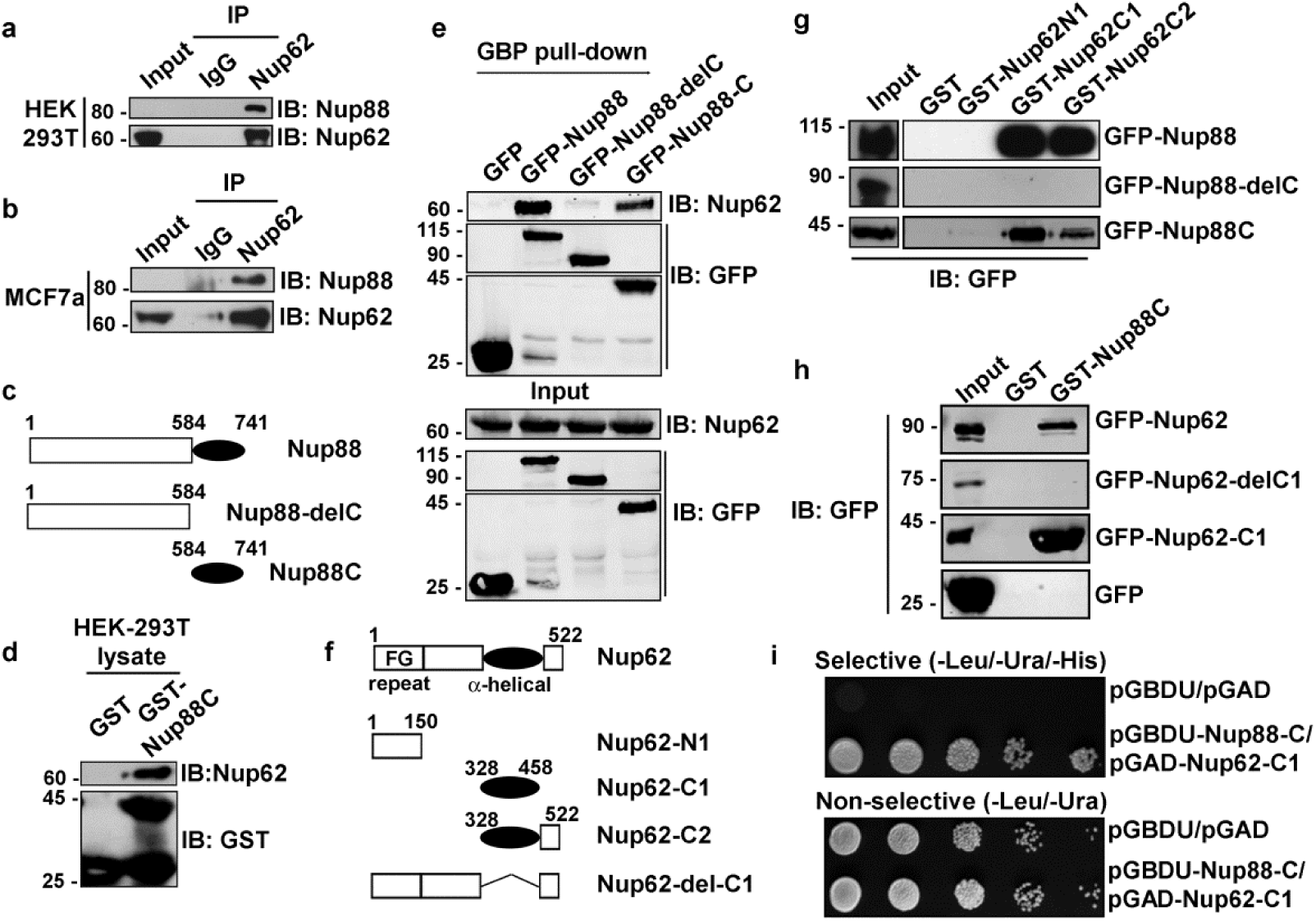
Nup88 and Nup62 interact through their carboxy-termini. (a) Immuno-precipitation (IP) using anti-Nup62 antibody from HEK293T cell lysates and detection of indicated proteins by western blotting. (b) Same as in (a) but from MCF7a cell lysates. (c) Schematic representation of Nup88 domain constructs used in cellular transfection and GST and GFP binding protein (GBP) pull-down experiments. GFP-Nup88 full length (1-741), GFP-Nup88delC (1-584), and GFP-Nup88C coiled-coil domain (585-741). (d) Pull down on GST and GST-Nup88C coated beads from HEK293T lysates and detection by immunoblotting (IB) with indicated antibodies. (e) GBP pull-down from HEK293T lysates expressing either GFP-Nup88 or GFP-Nup88delC or GFP-Nup88C. Pulldown and input samples were immunoblotted (IB) with the indicated antibodies. (f) Schematic representation of Nup62 constructs used in this study. Nup62 full length (aa 1-522), Nup62-N1 (aa 1-150), Nup62-C1 (aa 328-458), Nup62-C2 (aa 328-522), Nup62-del-C1 (Nup62 lacking aa 328-458). (g) Beads coated with proteins indicated on top of the lanes used in pulldown experiments from HEK293T cell lysates expressing GFP-Nup88, or Nup88delC or Nup88C proteins. IB of pulldown samples, and input fractions with the anti-GFP antibody. (h) Same as (g), but the HEK293T cell lysate is expressing GFP-Nup62 full length, or Nup62-del-C1 or Nup62-C1 or GFP. IB of pulldown samples and input fractions with the anti-GFP antibody. (i) Yeast two-hybrid interaction analysis using the DNA binding domain (pGBDU) construct of Nup88C and activation domain (pGAD) construct of Nup62-C1. Doubly transformed yeast colonies were grown on selective and non-selective media to score for the interaction. Images are a representative from at least n=3 repeat experiments.

### Nup88 and Nup62 interaction is cell cycle independent

Nucleoporins exhibit cell-cycle dependent differences in their subcellular localization and stability. The alpha-helical domain of Nup62 (Nup62-C1) assists in its centrosome localization ^31^. We asked how the localization of Nup88 and Nup62 changes during different cell-cycle phases. GFP tagged Nup88-fl, Nup88ΔC, Nup88C, Nup62-fl, Nup62ΔC1, and Nup62-C1 were generated to assess their subcellular localization (Fig. S4a, b). The Nup88-C did not localize to the nuclear envelope (NE), but the Nup62-C1 was found at the NE (Fig. S4b). We then asked if the NE localization of endogenous Nup88 changes under Nup62ΔC1 overexpression conditions. The Nup62-fl and Nup62-C1 reactivity were strong at the nuclear rim, but the Nup62ΔC1 remained diffused inside the nucleoplasm. Importantly, in all these cases, endogenous Nup88 was found in the NE (Fig. 4a, b, c), probably existing in a strong complex with endogenous Nup62 and Nup214. We asked if the Nup88-Nup62 interaction and their protein levels exhibit any cell cycle-dependent variations. Lysates from asynchronous HeLa cells, and once synchronized at G1/S, and mitotic phase (Fig. S5) were used in immunoprecipitation (IP) experiment using control (IgG) and anti-Nup62 antibodies. We detected the efficient IP of Nup88 under all cell synchronization conditions (Fig. 4d). Similarly, the Nup62-C1 also pulled out endogenous Nup88 (Fig. 4e), and Nup88-C pulled out endogenous Nup62 (Fig. 4f) from asynchronous and synchronized HeLa cell lysates. We next probed if the cell cycle stages affect levels of overexpressed Nup88. HeLa cells synchronized at G1/S boundary were released from the arrest for indicated time intervals. GFP-Nup88 was pulled down on GBP coated beads, and its interaction with Nup62 was determined. The interaction between Nup88 and Nup62 remained unperturbed (Fig. 4g, first panel). However, the GFP-Nup88 levels decreased ~18h post G1/S release, a time-point indicative of the early G1 phase (Fig. 4g, third panel). In contrast, the endogenous Nup62 levels did not change significantly (Fig. 4g, second and fourth panels). Our data highlights the fact that Nup88-Nup62 interaction is independent of cell-cycle, and Nup62 glycosylation status. Why the overexpressed Nup88 should be unstable in the early G1 phase is unclear.

**Fig. 4:**
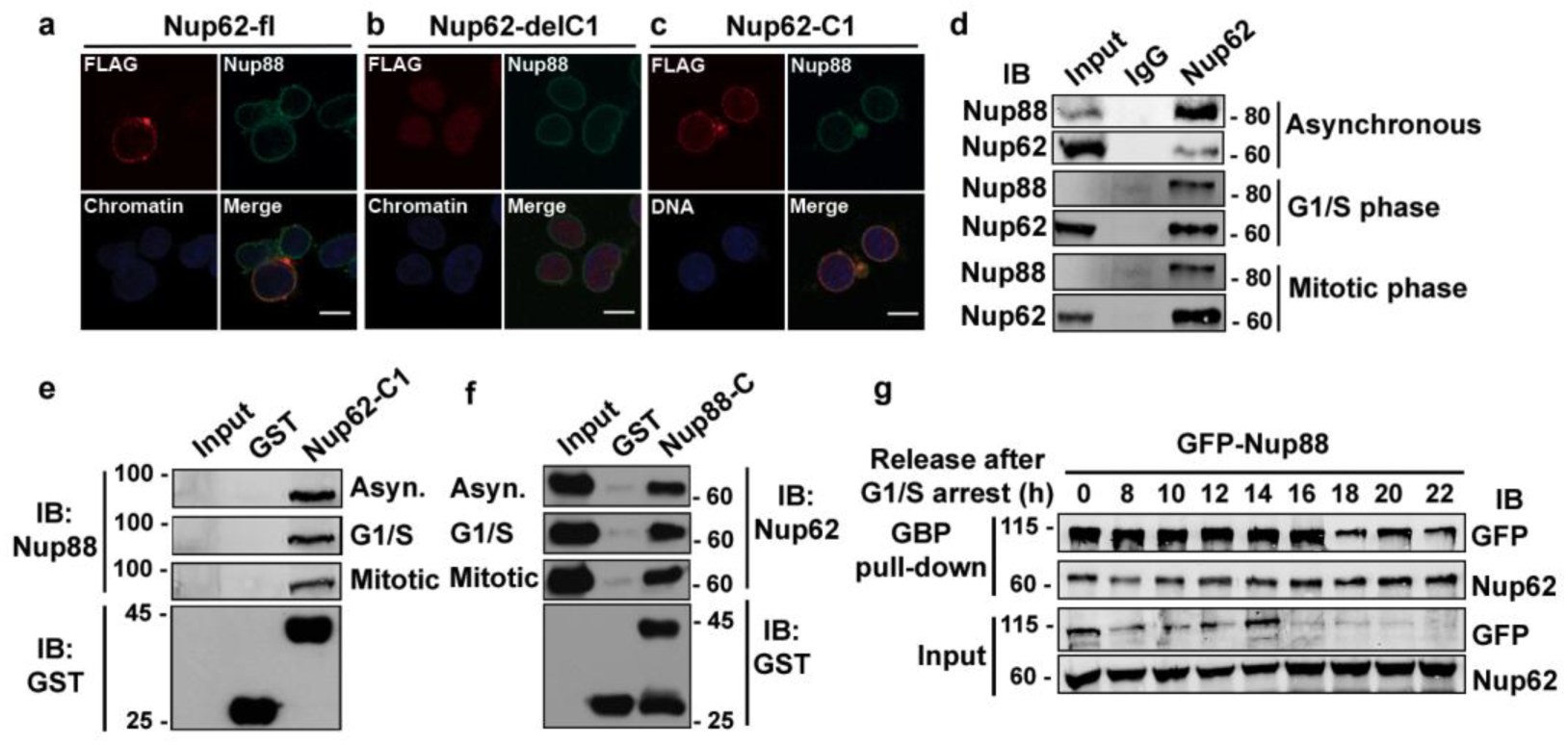
Nup88 interacts with Nup62 independent of the cell cycle phases. (a, b, c) Localization of FLAG-Nup62 constructs Nup62-fl, Nup62del-C1, and Nup62-C1 respectively in cells and detection by anti-FLAG (red) and anti-Nup88 (green). DNA is stained with Hoechst 33342. Scale bar =10µm. (d) Immunoprecipitation (IP) using control and anti-Nup62 IgG from HeLa lysates synchronized in different phases (as indicated) of the cell cycle. The IP sample was immunoblotted (IB) with anti-Nup88 and anti-Nup62 antibodies. (e) Pull down from HeLa lysates synchronized in indicated phases of cell cycle on beads coated with proteins GST or GST-Nup62-C1. The pull-down material is immunoblotted with the anti-Nup88 antibody. The GST tagged proteins were detected by anti-GST antibodies. (f) Same as in (e), but the beads are coated with GST or GST-Nup88C, and immunoblotting (IB) is performed with the anti-Nup62 antibody. (g) GBP pull down from GFP-Nup88 expressing HeLa cell lysates prepared from cells released for indicated time intervals after synchronization at G1/S phase. Pull down, and input fractions were immunoblotted (IB) with anti-GFP and anti-Nup62 antibodies. Images are a representative from at least n=3 repeat experiments.

### Nup62 interaction with Nup88 protects Nup88 from ubiquitination mediated degradation

The observation that overexpressed Nup88 is unstable, and the lack of Nup62 glycosylation degrades Nup88 ^18^ attributes strong stabilizing effects to Nup62 interaction. We probed the stabilizing effect of Nup62 on Nup88. GFP-Nup62 (present) or GFP (absent) expressing cells were treated with cycloheximide to prevent fresh protein synthesis, and levels of endogenous Nup88 were detected in total lysates. The endogenous levels of Nup88 were variable and often low and generally undetectable but expression of Nup62 seems to have stabilized Nup88 levels (Fig. S6a). When GFP-Nup88 overexpressing cells lost 50% of GFP-Nup88 within 2h of cycloheximide treatment, the endogenous Nup62 levels remained unchanged (Fig 5a). Moreover, distinct stabilization of GFP-Nup88 (from -/+ FLAG-Nup62, ~2 folds at t=0h to -/+ FLAG-Nup62 ~8 folds at t=4h) was observed when FLAG-Nup62 co-expressing cells were treated with cycloheximide (Fig. 5b). These observations indicate that overexpressed Nup88 is stabilized by co-expressing Nup62.

**Fig. 5:**
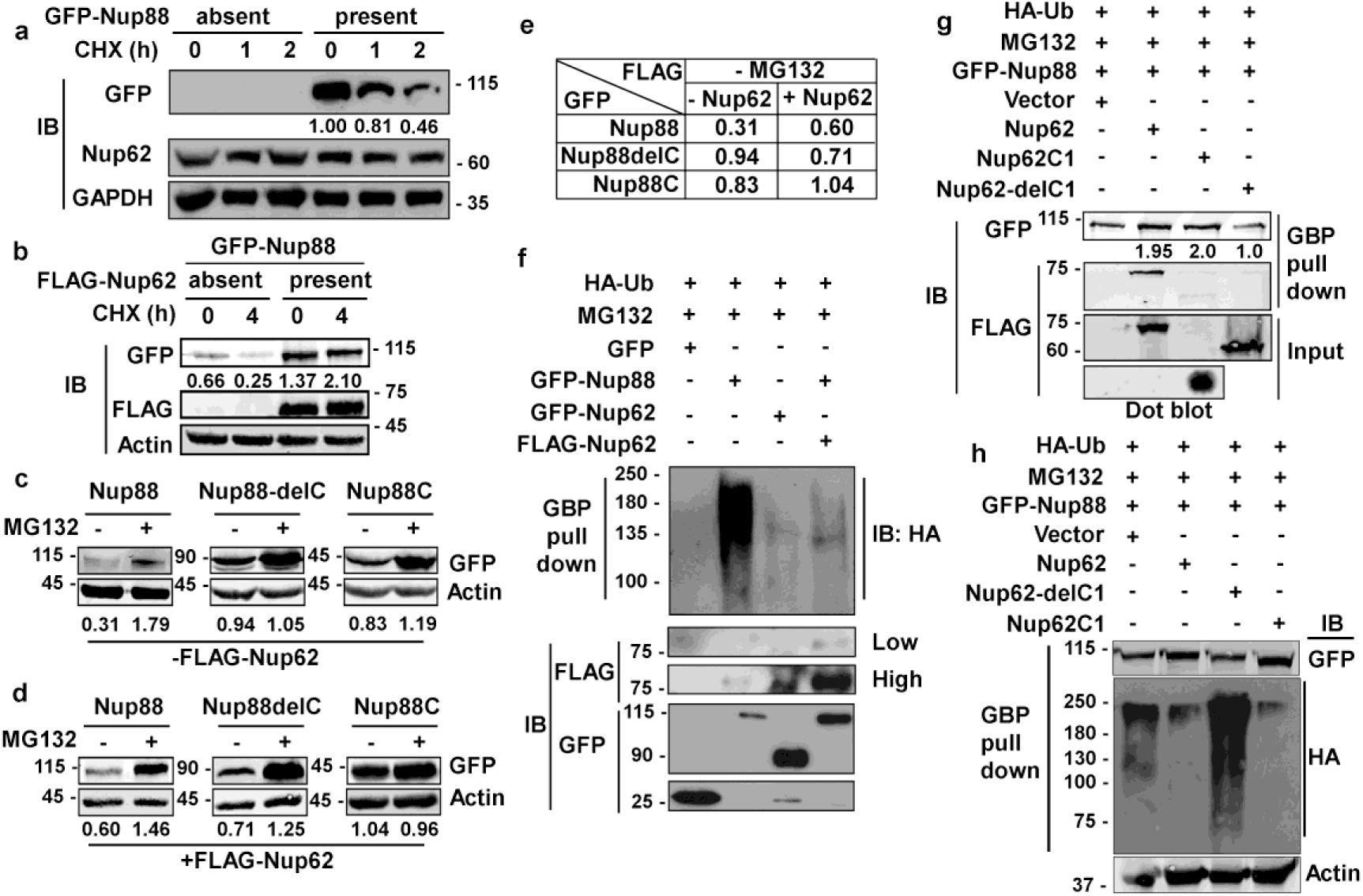
Nup62 stabilises Nup88 protein by protecting it from degradation. (a) Untransfected and GFP-Nup88 transfected cells were treated with cycloheximide for the indicated time points (h). Total lysate prepared from cells were immunoblotted with anti-GFP, anti-Nup62, and anti-GAPDH antibodies. (b) GFP-Nup88 transfected cells, cotransfected with mock (absent) or 3x-FLAG-Nup62 (present), were treated with cycloheximide for the indicated time points (h). Total cell lysates were immunoblotted (IB) with anti-GFP, anti-FLAG, and anti-Actin antibodies. (c) HEK293T cells transfected with Nup88 constructs indicated on top of panels were co-transfected with vector control (-FLAG-Nup62) or (d) FLAG-Nup62 (+FLAG-Nup62) and not-treated or treated with MG132. Lysates prepared from these cells were immunoblotted with anti-GFP and anti-Actin antibodies. (e) Table showing the stability of Nup88 constructs in the presence and absence of Nup62. (f) HEK293T cells were transfected with HA-Ubiquitin and treated with MG132. These cells were simultaneously co-transfected as indicated above the lanes. GBP pull-down was performed on the cell lysates, and pull down material was immunoblotted with anti-GFP, anti-HA and anti-FLAG antibodies. (g) HEK293T cells were transfected with HA-Ubiquitin and treated with MG132. These cells were simultaneously co-transfected as indicated above the lanes. GBP pull down material was probed with anti-FLAG, and anti-GFP antibodies. (h) HEK293T cells were transfected with HA-Ubiquitin and treated with MG132. These cells were simultaneously co-transfected as indicated above the lanes. GBP pull-down was performed on the cell lysates, and pull down material was immunoblotted with anti-GFP, anti-HA and anti-Actin antibodies. Images are a representative from at least n=3 repeat experiments.

We asked if the Nup88 degradation is ubiquitination dependent and if Nup62 interaction can provide stability against ubiquitination. Indeed, MG132 treatment increases endogenous Nup88 levels inside HeLa cells (Fig. S6b). In general, the presence of MG132 prevented degradation and accumulated Nup88 proteins. Interestingly, when the effect of Nup62 was assessed in the absence of proteasome inhibition, both the GFP-Nup88 (from 0.31 to 0.60) and GFP-Nup88C (from 0.83 to 1.04) were stabilized in the presence of Nup62. But, the Nup88ΔC construct lacking interaction with Nup62 was unstable (from 0.94 to 0.71) during Nup62 co-expression (Fig. 5c, d, e). We asked if the compromise in the stability of overexpressed Nup88 is due to ubiquitination. Cells overexpressing HA-Ubiquitin were treated with MG132, and GFP-Nup88 was expressed individually or in combination with Nup62. The samples overexpressing only GFP-Nup88 show strong ubiquitination, but the ubiquitin reactivity reduced to negligible levels when Nup62 was co-expressed (Fig. 5f). We next asked if the Nup62 interaction affects the degradation of overexpressed Nup88. Cells expressing HA-Ubiquitin and GFP-Nup88 were treated with MG132 and transfected with indicated FLAG-Nup62 constructs. GFP-Nup88 was pulled down on GBP beads followed with anti-GFP antibodies probing. We notice that the Nup62 and Nup62-C1 expression (capable of interacting with Nup88) impart more stability in Nup88 (~2 fold) than when Nup62ΔC1 (cannot interact with Nup88) is expressed (Fig. 5g). From similar experimental conditions, we established that Nup62 interaction is important for Nup88 stability. In cells, overexpressing GFP-Nup88 and HA-ubiquitin, Nup88 was ubiquitinated when Nup62ΔC1, a construct lacking the Nup88 interacting domain, was co-expressed. However, the co-expression of Nup88 interacting domain, Nup62C1, prevented Nup88 ubiquitination (Fig 5h).

### Nup88 interacts with NFκB and affects its downstream proliferative and inflammatory pathways

Nup88-214 sub-complex and Crm1 together regulate the nuclear export of cargo proteins like NFκB/*Dorsal* (p65) ^32^. In addition to genetic interaction, weak biochemical interaction between *Dorsal* and *mbo* (Nup88) is reported only from *Drosophila*. We asked if p65 and Nup88 can interact directly or the coiled-coil domains of Nup88 and or Nup62 play an important role in this interaction? First, we demonstrate that the tumor necrosis factor-α (TNF-α) induced normal nucleo-cytoplasmic distribution of p65 is unaltered when GFP-Nup88 is overexpressed (Fig. 6a). GFP-Nup88 pulldown showed efficient interaction with p65, but only in the presence of full length Nup62. Although the p65 interacted with Nup88 when Nup62-C1 was overexpressed but the strength of interaction was weak, and the same was absent with Nup62ΔC1 (Fig. 6b). We further tested the involvement of interacting domains, Nup88C and Nup62-C1, in p65 pulldown. Nup88C and Nup62-C1 pulled down endogenous Nup62 and Nup88, respectively, but both failed to individually pulldown p65 (Fig. 6c). Thus, the presence of stable Nup88 is imperative for interaction with p65. While analyzing the Nup88 expressing cells, we found overexpressed Nup88 inside the nucleus (Fig. S7a). Thus we asked if, in unstimulated cells, Nup88 can interact and sequester p65 inside the nucleus. Indeed the p65 was seen inside the nucleus of unstimulated GFP-Nup88 expressing cells (Fig. S7b). We further probed if nuclear p65 is active and can induce the transcription of its target genes. Comparative qRT-PCR analysis of p65 target genes in unstimulated GFP and GFP-Nup88 expressing cells suggest an increase in inflammatory cytokine, IL-6, levels (Fig. 6d) and of Ki-67 (an important proliferation antigen) levels (Fig. 6e). Further, enhancement in the expression of Akt and c-myc (growth and survival marker) and Bcl-2 and BIRC3 (apoptotic regulators) were seen (Fig. 6f). We strengthened the observation made in cell line overexpression studies by analysing p65 target genes in GEO & Oncomine oral cancer datasets already mentioned earlier for Nup88 and Nup62 upregulation. We found upregulation of IL6, Ki67, c-myc, Akt, and BIRC3 genes in the oral cancer dataset-GSE30784 (Fig 6g) as well as in analyzed head and neck statistics available at Oncomine (Fig. S8). Together, these observations indicate a direct interaction between Nup88 and p65, leading to the activation of the NFκB pathway during Nup88 overexpression.

**Fig. 6:**
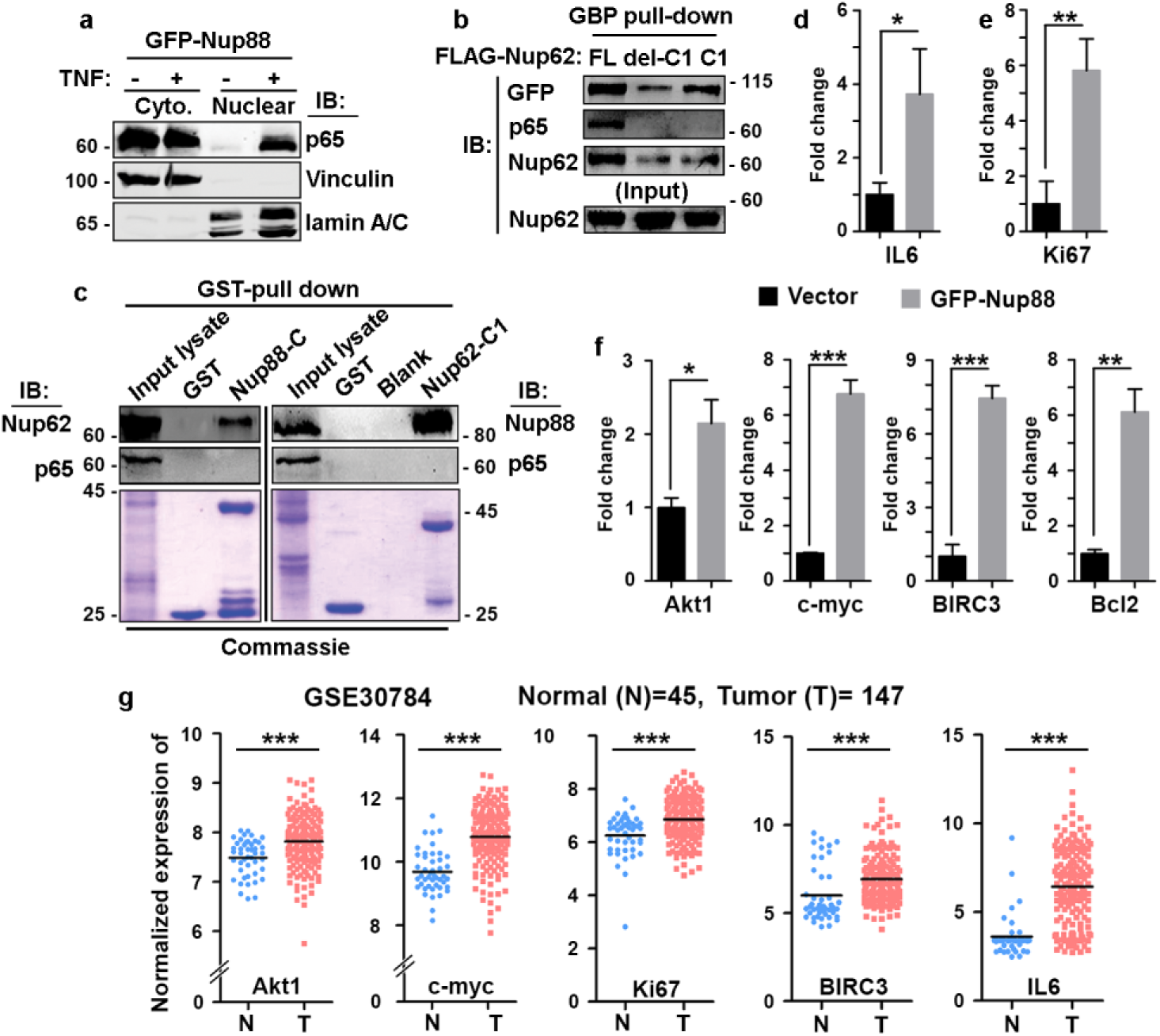
Stable Nup88 interacts with NFκB and activates downstream pathways. (a) Cytosolic and nuclear fractions prepared from GFP-Nup88 transfected cells left untreated (-) or treated (+) with tumor necrosis factor-α (TNF-α) were immunoblotted with anti-p65, anti-vinculin, and anti-lamin A/C antibodies. (b) GFP-Nup88 transfected cells were cotransfected with FLAG-Nup62 constructs indicated above the lanes. GBP pull-down material was probed with anti-GFP, ani-p65, and anti-Nup62 antibodies. (c) Beads coated with GST tagged proteins indicated on top of the wells were used in pulldown from HEK293T lysates. Pull down material was immunoblotted with anti-Nup62, anti-Nup88, and anti-p65 antibodies. Lower panels show Coomassie of input lysates and bead-bound samples. (d,e) Rps16 normalized qRT-PCR data for IL-6 and Ki67 from GFPNup88 transfected cells. (f) Actin normalized qRT-PCR data of indicated NFκB target genes from GFP-Nup88 transfected cells. (g) IL-6, Ki67, Akt, c-myc, and BIRC3 expression analysed through microarray data analysis of publically available oral cancer dataset (GSE30784) on GEO database. Error bars indicate mean values ± SEM. Asterisks indicate statistical significance (Student’s t-test) *P < 0.05, **P < 0.01 and ***P < 0.001.

## Discussion

Nucleoporins regulate nucleocytoplasmic transport of biomolecules and thus also regulate gene expression. Nup107 complex is critical for mRNA export ^33–35^, and the Nup88-Nup214 complex regulates the transport of NFκB and pre-ribosomal assemblies ^32,36–39^, and hence critical for transcription and translation processes. Nup88 overexpression is becoming synonymous with cancer progression ^6^. We find elevated Nup88 and Nup62 mRNA and protein levels in oral cancer tissues and a positive correlation between elevated Nup88 levels vis-a-vis poor survival rates (Fig. 1). In accordance, the TCGA and MiPanda analysis on oral cancer tissues showed a significant increase in Nup62 and Nup88 levels. The endogenous Nup88 is in a stable complex, and its functions are tightly regulated. Possibly the co-overexpression of Nup62 and Nup88 allows the formation of a stoichiometric complex stabilizing Nup88 manifesting cancerous outcomes. While Nup88 protein levels were reported to be increasing with progressive stages of cancer ^40^, overexpression of no other nucleoporin could parallel these phenotypes ^11^. Importantly, we observe a constant increase in Nup62 protein levels in oral cancers.

Cells overexpressing Nup88 were reported to induce multi-nuclear structures ^12^, although a clear understanding of a suitable mechanism lacked. Overexpression of Nup88 or Nup62 allowed increased cell proliferation, colony-forming abilities, and migration properties (Fig. 2). We argue through our data that the Nup62 overexpression phenotypes assessed in cell culture setup were most probably an outcome of stabilized Nup88. Nup62 glycosylation levels are a critical player of Nup88 stability ^18^. However, our *in vitro* data suggest that Nup62 glycosylation is dispensable for the coiled-coil domain interactions with Nup88 (Fig. 3). But, the possibility of Nup62 glycosylation being an important player for Nup88 stability in the cellular context is still exists.

Nucleoporins do exhibit a cell-cycle dependent difference in their stability and localization and thus are involved in cell-cycle specific interactions ^41^. Nup62 is not a stable member of the Nup88-214 subcomplex, but it displays cell-cycle stage independent stable interaction with Nup88 without affecting its nuclear envelope localization (Fig. 4). Only the constituent nucleoporins of a defined subcomplex, like that of the Nup107 complex, exhibit such cell-cycle independent interactions ^42^. Interestingly, overexpressed and endogenous Nup62 remains stable, but the overexpressed Nup88 degrades over time in the absence of fresh protein synthesis, which concurs with observation that Nup88 is ubiquitinated, but Nup62 is not. Proteasomal inhibition studies further suggest that enabling Nup62 interaction diminishes Nup88 ubiquitination and degradation (Fig. 5). Such stabilizing interaction is known for other proteins, including that of interaction between NFκB and its inhibitor κB. Thus co-overexpression of Nup62 in cancers may probably work through the stabilization of Nup88, and stable Nup88 can engage in proliferative activities inducing tumorigenic transformation.

*Drosophila* Nup88 (*mbo*) interacts with *dorsal* (p65) and perturbs p65 nuclear export when the immune pathway is activated ^36^. Crm1 null flies induce p65 accumulation inside the nucleus upon bacterial challenge ^37^. Another nucleoporin, ELYS, is reported to affect the intranuclear dynamics of p65 ^43^. Contrary to the previous reports ^36,44^, Nup88 interacted strongly with p65, and the Nup88-p65 interaction seems direct (Fig. 6). It is apparent that Nup62 is not mediating the Nup88-p65 interaction, and this could also mean that the stability of overexpressed Nup88 is the prime outcome of co-expressed Nup62 in cancers, including oral cancer. NFκB signaling plays an important role in cell proliferation, anti-apoptosis, and inflammation ^45^, and the inflammatory milieu is known to support tumor growth and progression ^46^. Nup88-p65 interaction induces the expression of these pro-growth and anti-apoptotic molecules to orchestrate a conducive environment for tumorigenesis. The upregulation of IL-6 in unstimulated GFP-Nup88 expressing cells is an unprecedented observation (Fig. 6). However, this observation aligns with the IL-6 expression and inflammatory milieu in cancer ^47^. We suggest that in cancers overexpressing Nup88 or Nup88 and Nup62, stabilized Nup88 perhaps deregulates the transcriptional paradigm, and a dysregulated Nup88-p65 axis is inclined towards p65 dependent inflammatory and pro-growth response. In agreement with a recent report suggesting multiple roles for overexpressed Nup88 ^25^, NFκB pathway activation seems one of the mechanisms operating in Nup88 overexpressing cancer (Fig. 7).

**Fig. 7:**
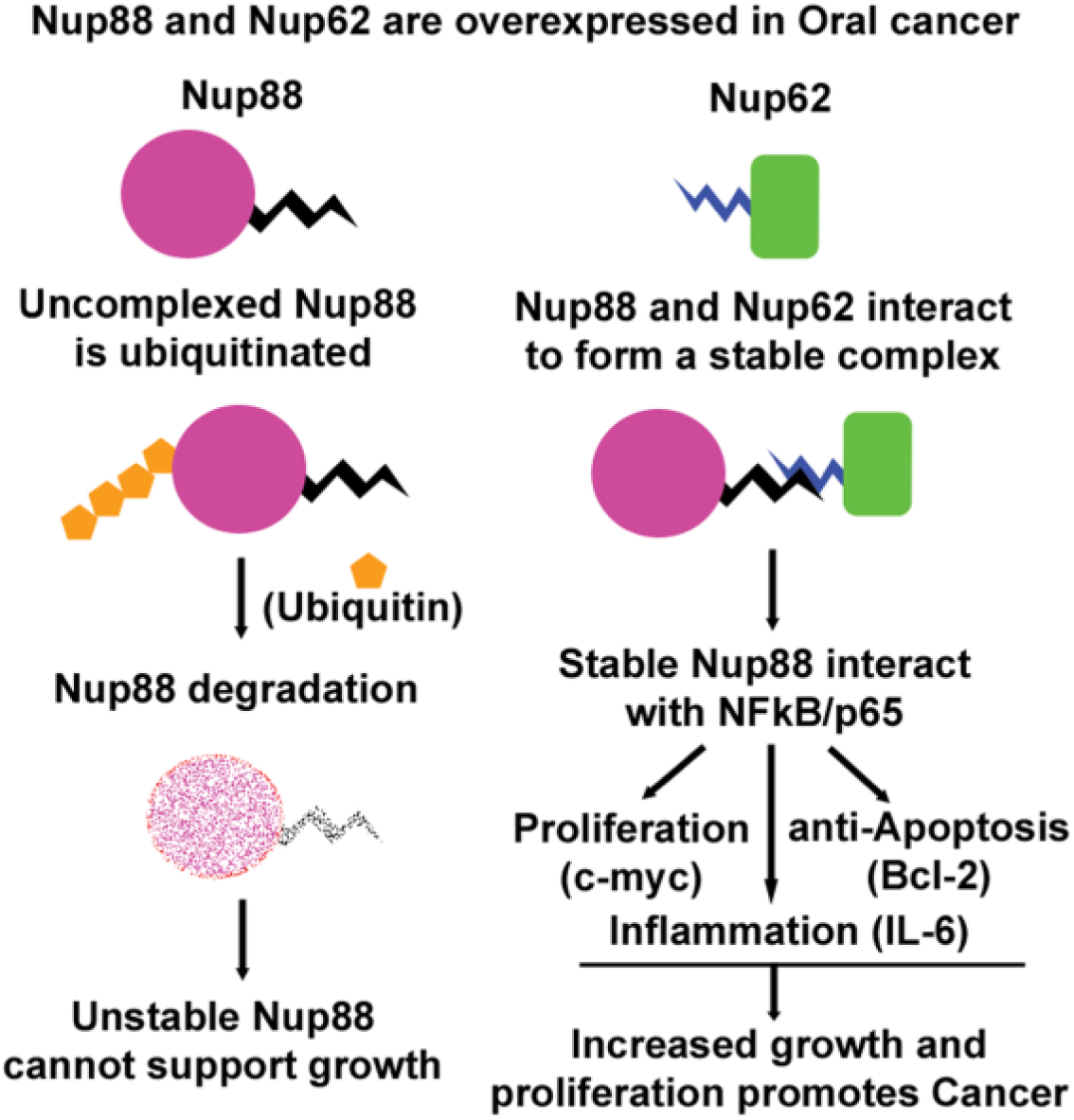
Stable Nup88 affects pro-growth and survival arm of NFκB pathway in cancer. Nup88 and Nup62 are overexpressed in oral cancer. Uncomplexed Nup88 is targeted for degradation by ubiquitin-mediated proteasomal pathway, whereas Nup88 complexed with Nup62 is stabilized. Stabilized Nup88 interacts with NFκB (p65) and promotes the expression of NFκB target genes-c-myc, Bcl2, and IL-6 to drive growth, anti-apoptosis, and inflammation simultaneously driving tumorigenesis.

In summary, we report the overexpression of Nup88 and Nup62 in oral cancer tissues. The conserved interaction between Nup88 and Nup62 primarily stabilizes Nup88 against ubiquitination mediated degradation. Overexpressed and stable Nup88 enriches partially in the nucleus and engages in strong interaction with NFκB. The unique presence of NFκB in the nucleus of unstimulated Nup88 overexpressing cells favours an inflammatory and anti-apoptotic survival environment to induce neoplastic transformations.

## Material and Methods

### Cancer tissue collection

Head and neck cancer tissue samples (T) and adjacent normal tissues (N) were collected from Bansal Hospital, Bhopal, India. The study was approved by the Institute Ethics Committee of Indian Institute of Science Education and Research (IISER) Bhopal and samples were collected with the consent of the patients. The tissues were snap-frozen immediately after surgery and stored at −80°C until use. The tissues for RNA isolation were collected in RNA Later (Thermo Fischer Scientific, AM7024). The clinical characteristics of patients used in the study are listed in Supplementary Table 5.

### Bioinformatics analysis

The co-expression plot for Nup88 and Nup62 in normal, primary cancer, metastasis and cell lines for oral cancer was extracted from MiPanda (http://mipanda.org). The cancer stage-specific expression was analyzed using Cancer RNA-Seq Nexus (CRN) (http://syslab4.nchu.edu.tw/). The differential expression graph for both the genes was plotted using GraphPad Prism. The survival curves specific to Nup88 and Nup62 were obtained from OncoLnc (http://www.oncolnc.org/). The differential expression pattern of Nup88 and Nup62 in oral cancer was analyzed with the help of Oncomine database. The graph of analyzed expression data of Nup88 and Nup62 in oral cancer was saved for the representation.

### Plasmids

The full-length construct of Ubiquitin (HA-Ubiquitin Plasmid #18712) and Nup88 (pEGFP-Nup88 Plasmid #64283) were obtained from Addgene. The C-terminal coiled-coil domain (Nup88-C, amino acids 585-741) of Nup88 was PCR amplified from human testis cDNA and cloned into pGEX6P1 with EcoRI and SalI enzyme sites. This domain was further subcloned into pEGFPC1. The Nup88 construct lacking the coiled coil domain (Nup88∆C) was created by inserting a stop codon after 584aa through site directed mutagenesis (Q5 Site Directed Mutagenesis Kit, NEB-E0554S) using pEGFPC1-Nup88 as template. pEGFPC1-Nup62 was a kind gift from Dr Radha Chauhan (NCCS, Pune). The N-terminal and C-terminal truncations of Nup62 were PCR amplified and cloned into pGEX6P1, pCMV3Tag1a and pEGFPN1 vectors. The yeast-two hybrid constructs for Nup88 and Nup62 truncations were made by subcloning them into pGADC1 and pBDUC1 vectors. Nup88-C (aa 585-741) and Nup62-C1 (aa 328-458) coding sequence were inserted into pGADC1 vector harbouring GAL4 activation domain (AD) and pBDUC1 vector harbouring GAL4 DNA binding domain (BD).

### Reagents and Antibodies

Reagents and antibodies used in this study were purchased from miscellaneous sources. Cycloheximide (Catalog no - 100183) was from MP Biomedicals and used at 1 µg/ml for different time points. MG132 (Catalog no - 474787-10MG) was from HiMedia and used at 10 µM concentration for 8h. The antibodies used for western blotting are anti-Nup88 (BD Biosciences# 611896, 1:2000 dilutions), anti-Nup62 (BD Biosciences# 610497, 1:6000 dilutions), anti-GAPDH (Abgenex# 10-10011, 1:6000 dilutions), anti-GFP (Santa Cruz# 9996 1:5000 dilutions), anti-GST (1:500 dilution), anti-HA (Sigma# H6908, 1:2000 dilutions), anti-FLAG (Sigma# F7425, 1:2000), anti-Actin (BD Biosciences# 612656, 1:5000), anti-Lamin A/C (BD Biosciences# 612162, 1:2000), anti-NFκB p65 (BD Biosciences# 610868, 1:2000), goat anti-rabbit IgG-HRP (GeNei# 114038001A, 1:10000), goat anti -mouse IgG-HRP (GeNei#, 114068001A, 1:10000).

### Cell Culture

HEK293T, HeLa and MCF-7a cells were obtained from American Type Culture Collection (ATCC). The cells were grown and maintained in Dulbecco’s Modified Eagle’s Medium (Gibco# 11995 supplemented with foetal bovine serum (Invitrogen# 16000044) and antibiotics (100 units/ml Penicillin and Streptomycin, Invitrogen# 15140122) in humidified incubator with 5% CO_2_ at 37°C.

### Cell synchronization and FACS analysis

HeLa cells were synchronized in G1/S phase by double thymidine block and into M-phase by a thymidine block followed by nocodazole treatment. Cells were cultured at 30% confluency, and two cycles of 2 mM thymidine was added for 18h with 9h post-release between the two treatments to synchronize cells at G1/S boundary. For the M-phase block, cells were cultured at 40% confluency and treated with 2 mM thymidine for 24h. Cells were then released for 5h and treated with 100 ng/ml nocodazole for 12h. Shake-off method was used to collect the mitotic cells. For FACS analysis, HeLa cells blocked in G1/S were trypsinized, and the cells blocked in M-phase were collected by mitotic shake-off, washed twice with PBS, and fixed in 70% ethanol at −20°C for 12h. The fixed cells were resuspended in PBS containing 50 µg/ml each of RNase A and propidium iodide. The cell cycle distribution was acquired by BD Calibur flow cytometry and analyzed by Modfit LT software.

### Lentivirus production

HEK293T cells were transfected with pLKO.1 shRNA plasmid (Sigma, Mission human genome shRNA library) and packaging plasmids-delta 8.9, VSV-G in a ratio of 10:5:1 with polyethylenimine (PEI) following the standard protocol. After 12h the media was replaced with fresh DMEM media containing 10% FBS and antibiotics. After 24h and 48h the supernatant was collected and spun to remove the cellular debris. The supernatant was filtered through a 0.45µm filter and stored at −80°C until further use.

### RNA interference and quantitative RT-PCR

MCF-7a cells were seeded in a six-well culture plate. After 12h incubation cells were infected with lentivirus containing gene-specific shRNA and control shRNA using 8 µg/ml polybrene containing media. The cells were selected with 0.8 µg/ml puromycin (Sigma, P9620) for 72h. Total RNA from cultured cells or tumor tissue was extracted by TRIzol (M.P. Biomed #15596018) method. The genomic DNA contamination was removed by RNAse free DNAse. 1 µg of RNA was reverse transcribed to cDNA by iScript cDNA synthesis kit (BioRad #17088) as per the manufacturer’s instructions. RT-PCR was performed using SYBR Green PCR master mix on a (BIO-RAD CFX Connect™ Real-Time System) Rps16 and Actin gene was used as the control gene, and the relative transcript level was calculated by CT value (2^−ΔΔCT^). Student’s t-test was used to compare the differences in the gene expression and p value < 0.05 was considered significant. The primers used are listed in the Supplementary Table 6.

### Cell fractionation

Cells were harvested after the respective treatment and washed twice with 1X PBS and resuspended in Extraction Buffer A (10 mM HEPES, 1 mM EDTA, 1 mM EGTA, 10 mM KCl, 1 mM DTT, 5 mM NaF, 1 mM Sodium vanadate, 10 mM Sodium molybdate, 0.5 mM PMSF and 1X Protease inhibitor cocktail). It was incubated on ice for 15 min. 0.3 µl of 10% NP-40 was added to it and vortexed for 30sec at 4°C. The lysate was centrifuged at 10000g for 1min at 4°C. The supernatant was collected as the cytosolic fraction. The pellet was resuspended in Extraction Buffer B (20 mM HEPES, 1 mM EDTA, 1 mM EGTA, 400 mM NaCl, 1 mM DTT, 5 mM NaF, 1 mM Sodium vanadate, 10 mM Sodium molybdate, 0.5 mM PMSF and 1X Protease inhibitor cocktail). It was incubated for 30 min on a shaker at 4°C and then centrifuged at 20000g for 5 min. The supernatant was collected as nuclear fractions.

### Western blot

HEK293T, MCF7a and HeLa cells were lysed in RIPA buffer (50 mM Tris pH 7.5, 10 mM EDTA, 1 mM EGTA, 150 mM NaCl, 1% Triton X 100, 0.2% Sodium deoxycholate and 1X Protease Inhibitor (Amresco M250). Cell lysate were sonicated and centrifuged at maximum speed (14800 rpm) to collect the supernatant. The homogenized head and neck tissues were lysed in GLyse AT buffer (GCC BIOTECH-GPA-004). The total protein is quantified using Bradford assay and samples were prepared by adding 6X SDS sample buffer and boiled for 10 min at 100°C. The electrophoresed protein samples were transferred to polyvinylidene difluoride membrane (PVDF) and blocked with 5% (w/v) non-fat milk and probed with suitable primary and secondary antibodies. The bands were detected with enhanced chemiluminescence substrate (BIO-RAD Clarity™ Western ECL Substrate Catalog no - 170-5060) method or by using the Odyssey infrared imaging system (LICOR Odyssey).

### Immunoprecipitation

The confluent HEK293T, MCF7a and HeLa cell monolayer was lysed in 500 µl of RIPA buffer, sonicated and centrifuged at maximum speed. The collected supernatant was incubated with 5 µg of anti-Nup62, and anti-mouse IgG and incubated at 4°C on rocker 12h. 20 µl of Protein-G sepharose beads were added to it and further incubated at 4°C on rocker for 4h. Bead bound samples were centrifuged and unbound fractions were collected separately, and beads were washed 4 times with chilled PBS. The eluted protein samples were processed with 6X SDS sample buffer, and samples were analyzed by western blotting as described earlier.

### GST pull-down assay

The GST and GST-fused proteins were purified from bacterial strains-*E. coli* BL21DE3 Star and Codon plus cells. 20 µg of each protein was allowed to bind glutathione beads for 1h at 4°C on a rocker. The unbound protein was removed and the beads were washed 4 times with wash buffer (20 mM Tris, 150 mM NaCl and 1 mM EDTA). The pull-down was performed by adding 500 µg of HEK293T or HeLa cell lysate (asynchronous or synchronous, depending on experiment) and allowed to bind for another 1h at 4°C on rocker. The unbound fraction was removed, and washes were given as above. The eluted protein was analysed by western blotting.

### GFP-trap pull downs

The experiment was performed as described previously ^48^. In brief, HEK293T cells were transfected with pEGFP-Nup88, pEGFPN1-Nup62, and their truncations. 48h post-transfection cells were harvested, lysed, sonicated and centrifuged. The supernatant fraction was incubated with GST-GBP protein at 4°C on the rocker for 12h. 20 µl of glutathione beads were added and incubated at 4°C on rocker for 4h. The further washing and elution steps are same as the GST pull-down experiment.

### Yeast two-hybrid assay

The pGADC1 and pGBDUC1 constructs of Nup88 and Nup62 truncations were co-transformed into yeast two-hybrid strain PJ69-4A. Double transformants were obtained on a non-selective (lacking Leu and Ura, double drop out) media. The 10 fold serial dilutions of equivalent numbers of transformants were spotted on non-selective (lacking Leu and Ura, double drop out) and selective media (lacking Leu, Ura, and His, triple drop out) and incubated at 30°C for three days for transformants to appear.

### Cell viability assay

MCF-7a cells were transfected with pEGFPC1, pEGFP-Nup88, and pEGFPN1-Nup62 in six-well culture plate. 24h post-transfection cells were harvested, and 5×10^3^ cells were seeded in each well of a 96 well culture plate in triplicates and allowed to grow for another 24h, 48h, and 72h. The cell growth was measured by conversion of MTT-tetrazolium salt to formazan crystal. 20 µl of MTT (2 mg/ml) was added to each well, incubated for 4h and the reaction was terminated by adding 100 µl of DMSO. Viability and cell proliferation were assessed by measuring the optical density at 570 nm in a plate reader (BioTek Eon,11-120-611).

Similarly, for knockdown, MCF7a cells were transfected with Nup88 shRNA, Nup62 shRNA, and control shRNA in a six-well culture plate. The cells were selected for 72h with 0.8 µg/ml puromycin. After selection, cells were processed as above for viability and proliferation activity measurements.

### Wound healing assay

MCF-7a cells were transfected with pEGFPC1, pEGFP-Nup88, and pEGFPN1-Nup62 in a six-well culture plate setup. At 90% confluence, a scratch was created with a 10 µl pipette tip. The cellular debris (dislodged cells) was removed by thorough PBS washing. The cells were imaged at 24h, 48h and 72h intervals on an inverted microscope (Leica Microsystems Model-DMIL LED Fluo). The wound closure rate in each case was measured from images using TScratch software ^49^.

### Colony-forming assay

MCF-7a cells were transfected with pEGFPC1, pEGFP-Nup88, and pEGFPN1-Nup62 in a six-well culture plate. After 24h of transfection, cells were harvested and seeded (2000 cells/well) in six-well culture plate and were allowed to grow for 15 days. The cells were washed with PBS and imaged on an inverted microscope. Further, cells were fixed in 3:1 ratio of methanol: acetic acid for 5 min. After fixation, cells were stained with 0.05% crystal violet in methanol for 15 min. The cells were washed again with distilled water and imaged (Leica Microsystems Model-DMIL LED Fluo) to determine the colony size and area. The number and area of the colonies was determined with ImageJ software and graph was plotted using GraphPad. The same was repeated with the cells transfected with Nup88 shRNA, Nup62 shRNA, and control shRNA in 30 mm plate after 72h of selection with puromycin.

### Soft Agar assay

MCF-7a cells were transfected with pEGFPC1, pEGFP-Nup88, and pEGFPN1-Nup62 in a six-well culture plate. After 24h of transfection, cells were harvested and seeded (1000 cells/well) in six-well culture plate coated with 1.5% agarose gel in DMEM complete media. The cells were allowed to grow for 7days. During growth 50% of media was carefully exchanged every other day. The colonies formed were imaged after 7days under inverted microscope (Leica Microsystems Model-DMIL LED Fluo).

### Immunofluorescence

HEK293T cells were grown in six-well culture plate with coverslips. The cells were washed with PBS and pre-extracted using PHEM buffer (60 mM PIPES, 20 mM HEPES, 10 mM EGTA, 0.2% Triton X-100 and 4 mM MgSO_4_) for 5 min at room temperature (RT). Pre-extracted cells were fixed with 4% paraformaldehyde for 15 min and permeabilized with rehydration buffer (10 mM Tris pH 7.4, 150 mM NaCl, 0.1% Triton X-100) for another 15 min at RT. The cells were blocked using 5% normal goat serum for 30 min and incubated with corresponding primary antibody at 4°C 12h. After three PBS washes, Alexa-fluor conjugated secondary antibody (Alexa Fluor 488-Invitrogen A11034, Alexa Fluor 568-Invitrogen A11031-1:800 dilution) was added to the cells and allowed to incubate for an hour at RT. Again, three washes of PBS were given, and nuclei were stained with Hoechst 33342 (Invitrogen, 1 mg/ml, 1:5000 dilutions). The coverslips were mounted on slides with VECTASHIELD mounting medium (H1000), and images were captured on a Zeiss LSM 780 Confocal Microscope and Olympus Confocal Laser Scanning Microscope-FY3000. All images were analyzed using ZEN (Zeiss) or Image J software.

### Statistical analysis

The statistical analysis was performed with GraphPad Prism5. Student’s t-test was used to calculate the significance value. In the bar graphs, differences between two groups were compared using an unpaired two-tailed Student’s t-test. In case of cancer tissues analysis paired two-tailed Student’s t-test was used to study the significance. The differences were considered statistically significant with \**P<0.05, ** P<0.01* and *\*\*\* P<0.001*.

## Abbreviations

NUP: Nucleoporin
NF-κB: Nuclear Factor Kappa-light-chain-enhancer of activated B cells
IκB: Inhibitor of NF-κB
Crm1: Chromosomal Maintenance 1
APC/C: Anaphase promoting complex/cyclosome
PLK1: Polo like kinase 1
ROCK1: Rho-associated coiled-coil containing protein kinase 1
GAPDH: Glyceraldehyde 3-phosphate dehydrogenase
CRN: Cancer RNA-Seq Nexus
MTT: 3-(4,5-dimethylthiazol-2-yl)-2,5-diphenyl tetrazolium bromide
GBP: GFP Binding protein
PPL: Periplakin
ELYS: Embryonic large molecule derived from yolk sac
IL-6: Interleukin-6

## Authors Contributions

U.S. designed and performed all experiments, analyzed and interpreted data. R.K.M. designed experiments and interpreted data. R.K.M. and U.S. wrote the manuscript. A.S. provided cancer patient’s tissues for the experiments.

## Acknowledgements

We thank Dr. Radha Chauhan (NCCS, Pune) for generously providing the pEGFPC1-Nup62 construct. We also thank Dr. Chandan Sahi, Dr. Sourav Datta, Dr. Himanshu Kumar, and Dr. Sanjeev Shukla (IISER, Bhopal) for providing reagents helpful in this study. We thank Nup and Sumo Biology Group for discussions and comments on the manuscript. We also thank FIST project SR/FST/LSI-643/2015 for the Olympus Confocal Laser Scanning Microscope-FY3000. This work has been supported by SERB grant # EMR/2016/001819 and intramural funding from the Department of Biological Sciences Indian Institute of Science Education and Research Bhopal and GATE fellowship to U.S.

## Conflict of Interest

The authors declare that they have no conflict of interest.

## Notes

### Competing Interest Statement

The authors have declared no competing interest.

