## Supporting Information for "Overexpressed Nup88 stabilized through interaction with Nup62 promotes NFκB dependent pathways in cancer"

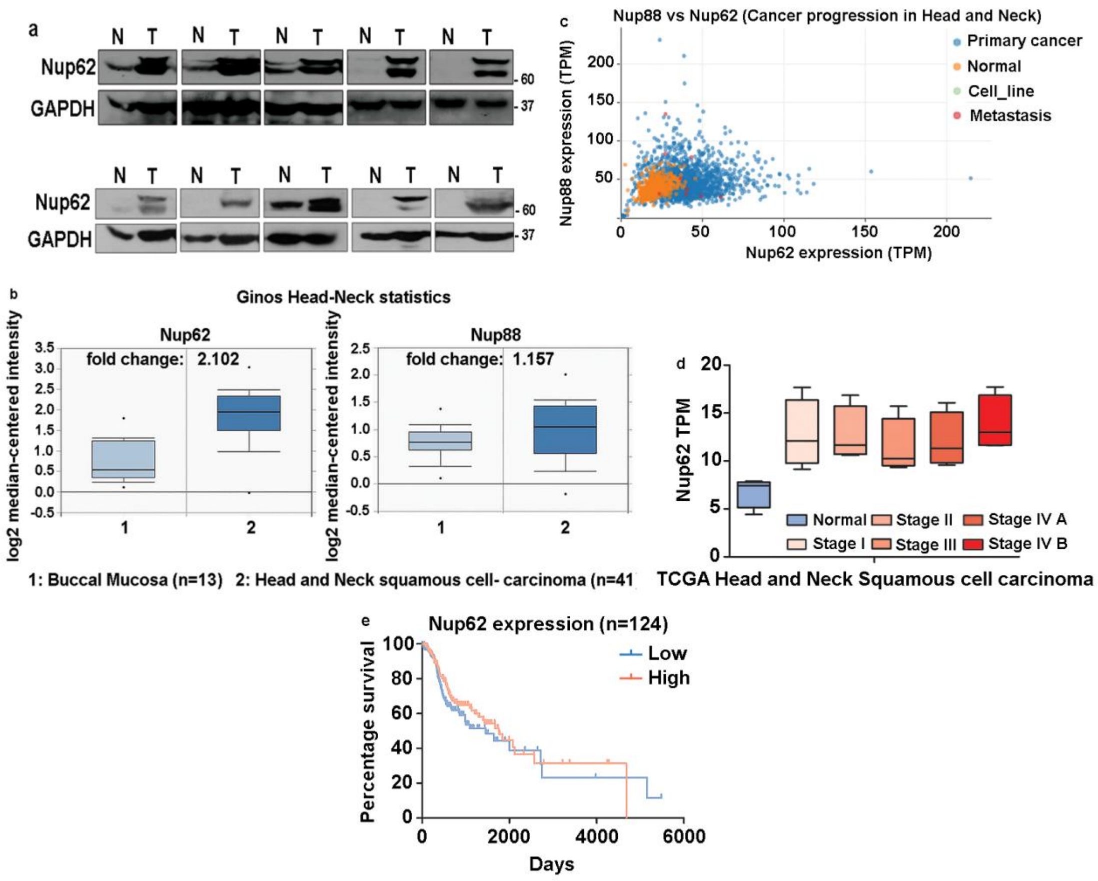


#### Fig. S1: Nup88 and Nup62 expression in head and neck cancer

(a) Western blot analysis of lysates prepared from oral cancer tissues using antibodies against Nup62, and GAPDH (N=Normal adjacent tissues, T=Tumor) (b) Nup62 and Nup88 expression analysis in head and neck cancer. Nup62 and Nup88 expression in Ginos head and neck statistics analyzed in Oncomine (*t–test*, $p<0.05$). (c) Co-expression analysis of available cancer datasets for Nup88 and Nup62. The plot is generated through Mi-Panda. (d) TCGA data for Nup62 TPM (Transcript Per Million) in different head and neck cancer stages analyzed using

Cancer RNA-Seq Nexus (*one-way analysis*, *p<0.05*). (e) Kaplan-Meier survival analysis of Nup62 expression using OncoLnc.

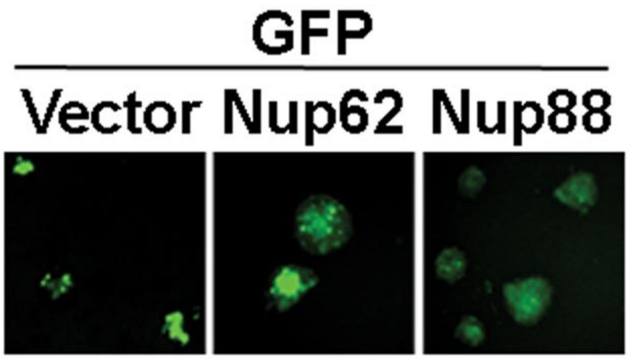


**Fig. S2: Expression of GFP, GFP-Nup62 and GFP-Nup88 constructs in colonies of MCF-7a cells**

MCF-7a cells transfected with GFP (Vector), GFP-Nup62 and GFP-Nup88 were imaged for GFP intensities in the soft agar assay using Fluorescent light microscopy.

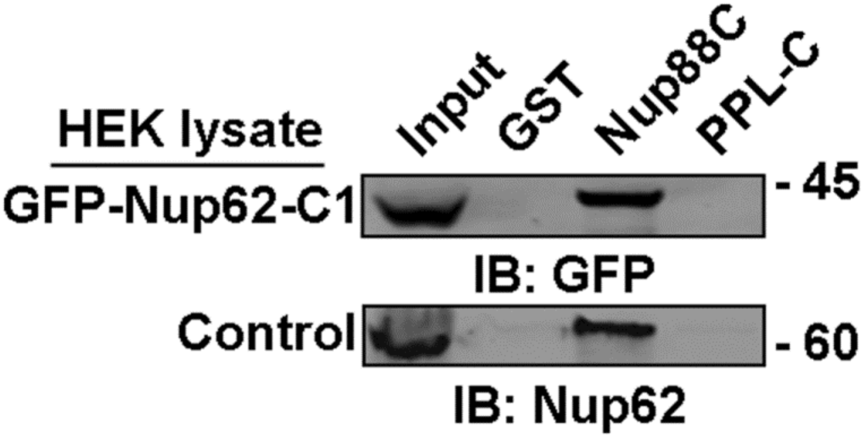


**Fig. S3: Nup88 coiled-coil interaction with Nup62 is specific**

Pulldown from control (untransfected) and GFP-Nup62C1 transfected HEK293T cell lysates on GSH beads coated with recombinant GST-tagged proteins as indicated on top of the wells each lane. Immunoblotting of pulldown material anti-GFP and anti-Nup62 antibodies.

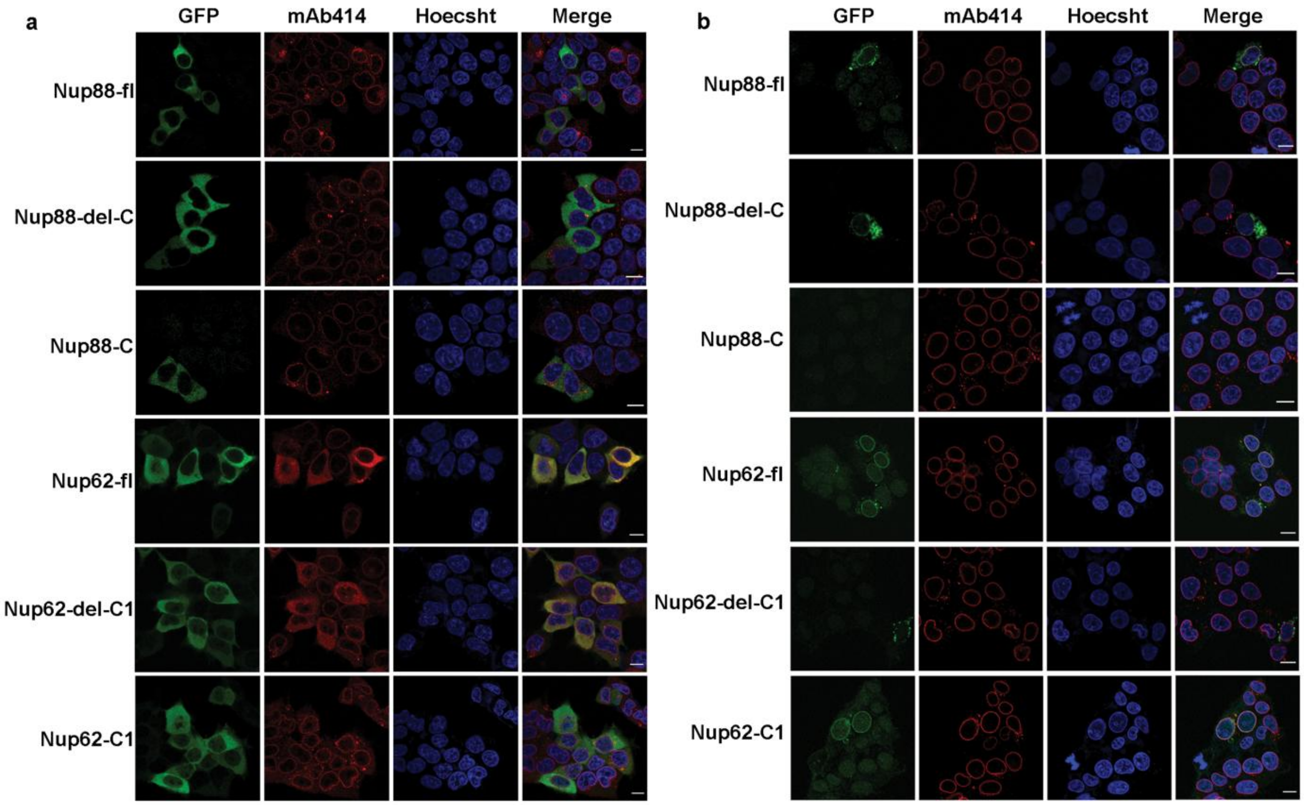


### Fig. S4: Localization of Nup88 and Nup62 constructs in cells

HeLa cells transfected with the indicated GFP-tagged Nup88 constructs- (Nup88-fl, Nup88delC, Nup88C) or Nup62 constructs - (Nup62-fl, Nup62delC, Nup62-C1). Cells were either not treated (non-pre-extracted, (a)), or treated (pre-extracted, (b)) with Triton-X-100 prior to fixation. Immunostaining with anti-GFP (green) and FG-Nup recognizing mAb414 (red) antibodies. DNA is stained with Hoechst 33342. Scale bar =10µm.

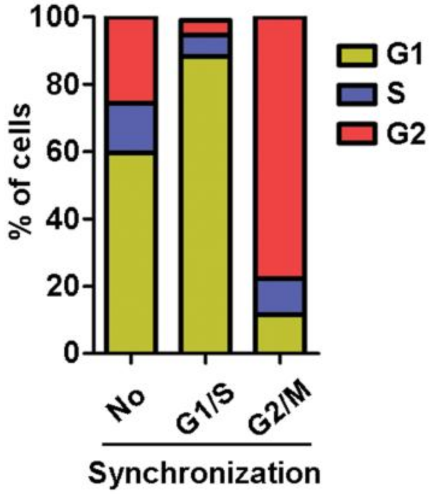


**Fig. S5: Synchronization of cells**

Graphical representation of FACS data under different treatments indicating the percentage of cells synchronized in different phases of the cell cycle.

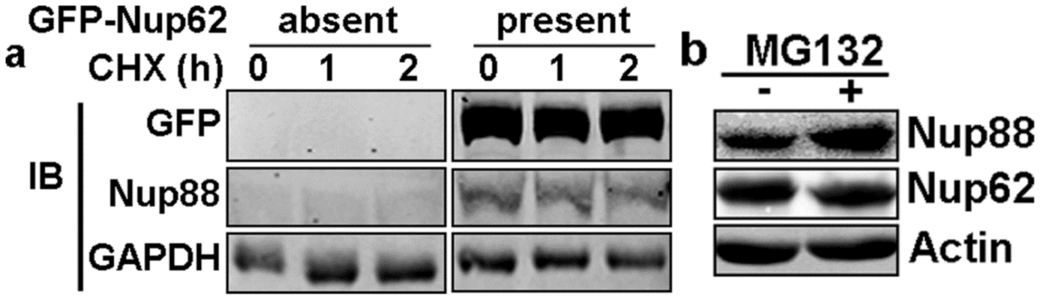


**Fig. S6: Nup88 is stabilized in presence of Nup62**

(a) GFP-Nup62 transfected (present) or GFP transfected (absent) cells were treated with cycloheximide for the indicated time points. Total lysate prepared from GFP transfected cells (left panels) and GFP-Nup62 transfected cells (right panels) were immunoblotted with anti-GFP, anti-Nup88, and anti-GAPDH antibodies. (b) Immunoblotting of lysates obtained from cells treated with (+) or without (-) MG132 with anti-Nup88, anti-Nup62, and anti-Actin antibodies.

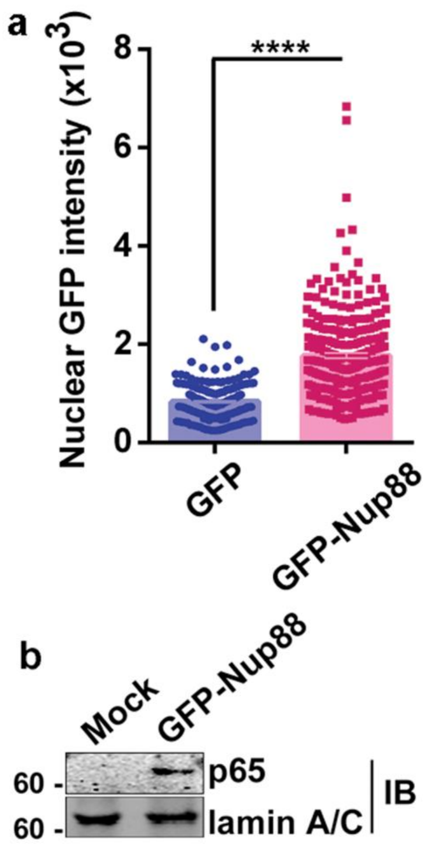


**Fig. S7: Nuclear localisation of Nup88 sequester p65 into the nucleus**

(a) Quantification of nuclear GFP signal intensities from GFP and GFP-Nup88 transfected HeLa cells using ImageJ Software. The graphs were plotted in GraphPad Prism6 and statistical significance was calculated from Student’s t-test. Asterisk represents the significance value, *****P* <0.0001 (b) Nuclear fractions from control and GFP-Nup88 transfected cells immunoblotted with anti-p65 and anti-Lamin A/C.

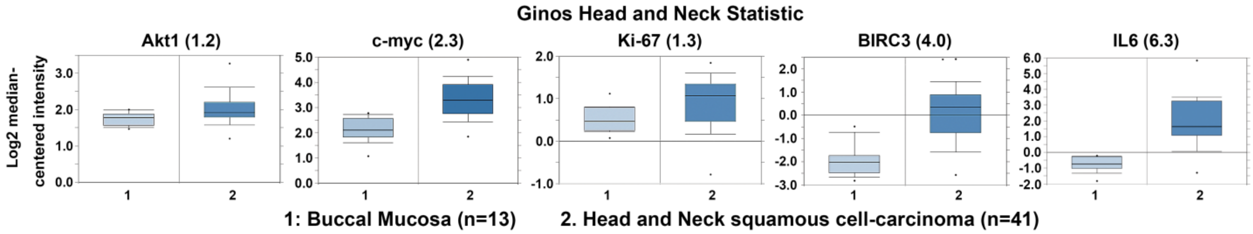


**Fig. S8: p65 target genes analyzed in Ginos Head and Neck Statistics at Oncomine.**

IL-6, Ki67, Akt, c-myc, and BIRC3 expression analyzed at Oncomine. The boxplots were downloaded from Oncomine for representation. The fold change for each gene is mentioned in the bracket.

**Supplementary Table**

Tables indicating area and the number of colonies obtained after overexpression of control, Nup62, or Nup88 in the colony-forming (Tables 1) and soft agar assay (Tables 3). Tables indicating area and number of colonies obtained after shRNA mediated knockdown of control, Nup62, or Nup88 in the colony-forming (Tables 2) and soft agar assay (Tables 4). The Table 5 represents the clinical characteristics of oral cancer patients and Table 6 enlists the primers used for qRT-PCR.

**Table 1**

Colony formation assay (GFP)

| Sample | Area of colony (mean ± SEM) | Number of colonies |
|---|---|---|
| Control | 37005 ± 3665 | N=43 |
| Nup62 | 51652 ± 4867 | N=41 |
| Nup88 | 57324 ± 6692 | N=46 |

**Table 2**

Colony formation assay (shRNA)

| Sample | Area of colony (mean ± SEM) | Number of colonies |
|---|---|---|
| Control | 71311 ± 4785 | N=110 |
| Nup62 | 54080 ± 6732 | N=42 |
| Nup88 | 22052 ± 3656 | N=13 |

**Table 3**

Soft Agar assay (GFP)

| Sample | Area of colony (mean ± SEM) | Number of colonies |
|---|---|---|
| Control | 2519 ± 209.3 | N=66 |
| Nup62 | 3986 ± 399.9 | N=109 |
| Nup88 | 3648 ± 320.0 | N=90 |

**Table 4**

Soft Agar assay (shRNA)

| Sample | Area of colony (mean ± SEM) | Number of colonies |
|---|---|---|
| Control | 2819 ± 254.3 | N=46 |
| Nup62 | 1952 ± 170.5 | N=47 |
| Nup88 | 1571 ± 147.5 | N=17 |

**Supplementary Table 5**: Clinicopathological characteristics of head and neck cancer patients.

| Sample ID | Gender | Age of Patient | Site of Cancer | Histopathology |
|---|---|---|---|---|
| S-1 | Male | 49 | Lower Lip | Squamous Cell Carcinoma (SCC) |
| S-2 | Male | 29 | Buccal Mucosa | Squamous Cell Carcinoma (SCC) |
| S-3 | Female | 60 | Lower GBS | Squamous Cell Carcinoma (SCC) |
| S-4 | Female | 46 | Buccal Mucosa | Squamous Cell Carcinoma (SCC) |
| S-5 | Male | 45 | Buccal Mucosa | Squamous Cell Carcinoma (SCC) |
| S-6 | Male | 55 | Buccal Mucosa | Squamous Cell Carcinoma (SCC) |
| S-7 | Male | 61 | Tongue | Coagulative necrosis |
| S-8 | Female | 58 | Tongue | Squamous Cell Carcinoma (SCC) |
| S-9 | Male | 60 | Buccal Mucosa | Squamous Cell Carcinoma (SCC) |
| S-10 | Male | 47 | GBS | Squamous Cell Carcinoma (SCC) |
| S-11 | Male | 70 | Buccal Mucosa | Verrucous Carcinoma |
| S-12 | Male | 53 | Buccal Mucosa | Squamous Cell Carcinoma (SCC) |
| S-13 | Male | 48 | GBS | Squamous Cell Carcinoma (SCC) |
| S-14 | Male | 42 | Oral Cavity | Squamous Cell Carcinoma (SCC) |
| S-15 | Male | 42 | Buccal Mucosa | Verrucous Carcinoma |
| S-16 | Male | 39 | Tongue | Squamous Cell Carcinoma (SCC) |
| S-17 | Female | 38 | Buccal Mucosa | Squamous Cell Carcinoma (SCC) |
| S-18 | Male | 48 | Tongue | Squamous Cell Carcinoma (SCC) |

**Supplementary Table 6:** List of primer with sequences
Quantitative real-time PCR primers (q-RTPCR)

| Serial No. | Gene ID | 5'-oligo Sequence-3' |
|---|---|---|
| 1. | Nup88 F | CTGCTTTGTACTCGAGAAGAT |
| 2. | Nup88 R | GCTGAGGAGCTGAAGGCATTCTTC |
| 3. | Nup62 F | CTACAAGCTGGCTGAGAACA |
| 4. | Nup62 R | ATGTGCGCATTGAGGATCT |
| 5. | Rps16 F | CGCGGCAATGGTCTCATCAAG |
| 6. | Rps16 R | GGAGATGGACTGACGGATAGCA |
| 7. | MKi67 F | AGAGTCAGGTTCAGAAATCC |
| 8. | MKi67 R | TCTTTCTCCCTCCTCTCTT |
| 9. | IL6 F | TACATCCTCGACGGCATCTC |
| 10. | IL6 R | CCAGGCAAGTCTCCTCATTG |
| 11. | c-Myc F | CTGAGGAGGAACAAGAAGATGAG |
| 12. | c-Myc R | TAGTTGTGCTGATGTGTGGAG |
| 13. | Akt F | CAAGGACGGGCACATTAAGA |
| 14. | Akt R | CCGCACATCATCTCGTACAT |
| 15. | BIRC3 F | TTTCCGTGGCTCTTATTCAAACT |
| 16. | BIRC3_R | GCACAGTGGTAGGAACTTCTCAT |
| 17. | Bcl2 F | GGTGGGGTCATGTGTGTGG |
| 18. | Bcl2 R | CGGTTCAGGTACTCAGTCATCC |
| 19. | Actin F | AACTGGGACGACATGGAGAAAA |
| 20. | Actin R | GGATAGCACAGCCTGGATAGCA |
